# Biomechanical stress in the retinal pigment epithelium surrounding laser scars in humans

**DOI:** 10.64898/2026.10.27.762316

**Authors:** Ysé Borella, Céline Chaumette, Jean-François Girmens, Cedric Dulière, Florence Rossant, Kiyoko Gocho, Daniel Lee, Ethan A. Rossi, José-Alain Sahel, Luis Almeida, Marie Darche, Michel Paques

## Abstract

**Purpose:** The wound response of an epithelium comprises tissue-scale activation of stress fibers whose contraction regionally reconfigures the epithelium and drives closure of the wound. Recent *in vitro* studies have shown that similar biomechanical processes can paradoxically lead to enlargement of epithelial holes. In patients having undergone laser photocoagulation of the retinal pigment epithelium (RPE), laser scars may expand over years. Here, to document the implication of epithelial biomechanics in the enlargement of laser scars we analyzed the dynamic changes affecting the RPE around scars in patients and performed confocal microscopy on a RPE flat-mount from a donor eye.

**Methods:** We analyzed the image files of five patients (age 61 to 85 years) showing enlargement of laser scars in the RPE (follow-up, 13 to 60 months). In three of them, prospective imaging was performed using transscleral illumination adaptive optics ophthalmoscopy (AOO), an imaging technology enabling visualization of RPE cells. Time-lapse sequences were analyzed to track dynamic processes in the RPE surrounding scars. A flat-mounted phalloidin-labeled RPE from a human donor eye with a history of scatter photocoagulation was examined by confocal microscopy to document the distribution of stress fibers and cell metrics.

**Results:** Time-lapse sequences showed that enlargement of laser scars was accompanied by the displacement of microscopic features in margins of atrophy. AOO imaging suggested locally coordinated displacement of mosaic of darks spots, with a generally radial or tangent direction relative to the edges of scars. Such movements were detected up to several hundred microns from the edges of scars. On the histology sample, elongated RPE cells with stress fibers forming actin cables were found along edges of scars, which formed fascicles between adjacent scars.

**Conclusion:** Enlargement of RPE scars may be accompanied by long-lasting, large-scale changes in the distribution of microscopic features in the margins of lesions. In particular, transscleral illumination AOO showed coordinated displacement of a mosaic similar to the one described for the RPE. The finding by histology of RPE cell with a tractional phenotype along edges further suggests that epithelial biomechanics contribute to such movements and may hence contribute to scar enlargement.

---

In post-mitotic epithelial monolayers, the response to a wound crucially relies on the activation of collective biomechanics [1]. Epithelial cells are interconnected through apical homodimeric cadherins junctions that transmit forces generated by intracellular actin-myosin complexes. *In vitro*, when challenged by a wound, epithelial cells reorganize actin-myosin complexes to form stress fibers connecting several cells, that may express a multicellular polarized expression of actin along the edges of the scar, so-called actin cable. The contraction of this actin cable regularizes the edge of wounds and close the gap through a purse-string effect, deforming and displacing cells toward the center of the scar [2–4]. The movements and shape of cells thus gives indications on the force vectors to which they are submitted [5]. The collective effect of cellular tractions can drive *en bloc* displacements of epithelial cells over millimetric distances [6]. These processes are phylogenetically conserved, being reported in epithelia from vertebrates and invertebrates*, in vitro* and during development [7–9]. Recently, in the context of a depleted cell density *in vitro*, cell contractility has been shown to paradoxically promote the expansion of epithelial holes [10]. This ambivalent effect of cell contractility suggests that there is a balance of inward versus outward forces, the net macroscopic (i.e. tissue-scale) effect depending on several factors among which is cell density and adhesion to the substrate.

The RPE is a monolayer of post-mitotic cells supporting critical functions for photoreceptors [11]. Laser photocoagulation, a common treatment for retinal vascular diseases, causes focal destruction of RPE cells. The ensuing scar may secondarily enlarge [12–14] and subsequently cause expanding scotoma over years. The mechanism driving expansion of scars is unknown. Propagation of photoreceptor necrosis or choriocapillaris occlusion have been suggested as possible mechanisms [15]. The clinical similarities of scar expansion with geographic atrophy (GA), the dry form of age-related macular degeneration, also suggests a role for degenerative mechanisms [16,17]. However, to our knowledge, there is no histology data to support the implication of these pathways during the expansion of RPE scar.

The present study originated from the clinical observations of patients in which comparison of fundus images revealed that expansion of laser scars was accompanied by displacement of periatrophic features such as hyperpigmented spots. This prompted us to investigate the possible implication of epithelial biomechanics in such mobilization. For this we analyzed the dynamics of the RPE both at the mesoscopic and microscopic scales and the spectrum of RPE cell shapes and actin expression by confocal microscopy of a donor eye.

## Methods

All procedures were performed according to institutional clinical studies carried out according to the principles outlined in the Declaration of Helsinki and approved by an ethics committee (Comité de Protection des Personnes Sud-Est III; registered in ClinicalTrial.Gov NCT041 28150 and NCT04129021). We searched our clinical database to identify patients having imaging files that fulfilled three criteria: expanding laser scars in the posterior pole, follow-up of at least one year, and image quality enabling to track features of the RPE. This led to the identification of five patients who were then proposed to participate in this study. All participants gave informed, written consent prior to inclusion. Their clinical characteristics are summarized in table 1. None had evidence of age-related macular degeneration in the included or fellow eye. Conventional color fundus photographs and scanning laser ophthalmoscopy (SLO; Spectralis system (Heidelberg Engineering, Heidelberg, Germany) were retrieved from image files. In addition, cases 3, 4 and 5 underwent prospective imaging using flood-illumination adaptive optics ophthalmoscopy (AOO) with transscleral fundus illumination (TFI) mode (ImagineEye, Orsay, France). This commercially available system uses two LED arrays at 810 nm which shine light to the fundus through the limbus; the light reflected from the sclera is detected through the pupil. This system has proven its capability to document the RPE mosaic [18,19].

**Table 1.** Clinical characteristics of patients (BRVO: branch retinal vein occlusion).

| Case | Age/sex | Visual acuity | Underlying disease | Follow-up (months) | Number of images (mean interval in months) | Growth rate of atrophy |
|---|---|---|---|---|---|---|
| 1 | 61/m | 20/20 | Diabetic retinopathy | 44 | 6 (7.3) | 136 µm/y |
| 2 | 85/m | 20/30 | Diabetic retinopathy | 60 | 14 (4.3) | 50 µm/y |
| 3 | 71/m | 20/40 | BRVO | 16 | 6 (2.7) | 36 µm/y |
| 4 | 75/m | 20/200 | BRVO | 14 | 4 (3.5) | 92 µm/y |
| 5 | 78/m | 20/25 | BRVO | 13 | 5 (2.6) | 24 µm/y |

TFI AOO images were montaged using i2k software (Imagine Eyes, Orsay, France, https://www.imagine-eyes.com/products/i2kretina). Time-lapse sequences of color fundus photographs, SLO and/or TFI AOO were constructed with Fiji 1.54P (developed by Wayne Rasband; available at rsb.inf.nih.gov/ij; National Institutes of Health, Bethesda, MD). Accuracy of alignments were visually evaluated based on anatomical landmarks such as vessels. Residual jitter was corrected using i2k Align and/or by manual correction using Adobe Photoshop 7.0 (Adobe Corporation, Mountain View, CA). Then, by scrolling time-lapse sequences at various frame rates (typically 5 - 10fps), changes in the distribution of fundus features over time and space were analyzed. The magnification factors provided by each system were used for the calculation of the expansion rate of atrophy, which was measured along the long axis of lesions. Appropriate plugins from Fiji were used to build up color coded maps and merged images. On time-lapse sequences, microscopic darks spots that could be tracked on at least three successive images were identified and their pathway and direction was reported on a directional map. Due to the faint contrast of image and their size, the length of the displacement was not considered reliable and was not reported.

For histology, one eye from an 84-year-old female donor with a history of branch retinal vein occlusion that had scatter laser photocoagulation was obtained from Lion’s Gift of Sight (Minnesota, USA) operating under the regulations of the U.S. Food and Drug Administration and the Eye Bank Association of America. Harvesting of the eye and fixation in paraformaldehyde 4% were done less than 24 hours post-mortem. Importation in France was done under relevant regulations for transfer of human tissues (CODECOH DC-2015-2400). Upon arrival in our lab, post-mortem *ex vivo* near infrared autofluorescence (NIRAF) imaging images using the Spectralis SLO was done to confirm the diagnosis and document the morphology of the scars. After careful ablation of the retina, the sample then underwent blocking and permeabilization in phosphate buffered saline (PBS) with gelatine 0.2% (PBSGT) (Prolabo, 24350262) and Triton X-100 0.5% (Merck Millipore Sigma-Aldrich, X-100) during 3 hours at room temperature. Then, sample was incubated with phalloidin-fluor 647 (ThermoFisher Scientific Invitrogen, A-22287) at 1:200 and DAPI (ThermoFisher Scientific Invitrogen) at 1:500, prepared in PBSGT then filtered with 22 µm filters for 2 hours. The sample was then incubated in TrueBlackPlus Lipofuscin Autofluorescence Quencher (Biotium, 23014) at 1:40 dilution 0.2% and triton X-100 5% in PBSGT for 20 minutes. Flatmount was mounted for confocal imaging in Rapid Clear 1.49 (SUNJin Lab, RC14900) medium. Confocal imaging was done with a Gasp Olympus microscope equipped with a red/far red spectral detector cube with 20x oil-immersion objective. A field of 4.5mm x 3.5mm centered on the scar was scanned by the confocal microscope. Images were montaged and analyzed by Imaris software (v9.5 to 9.7; Oxford Instruments, United Kingdom). Segmentation of the RPE cells on sampled macular fields of view (field of view size was 1.5×1.0mm for the three eyes (2 from 2 control patients (20681 cells in total), and 1 for the laser scar eye (2918 cells); selection of the field of view was based on the presence of a laser scar and the required microscope image quality for good segmentation) was done with the assistance of Cellpose, an open-source software using neural networks[20]. We used a custom model previously trained by us with data samples manually segmented and corrected for segmentation of RPE cells in dry AMD (5 samples of approximately 300 cells; cell diameter set to 45 pixels, flow threshold 0.4, cellprob and stitch thresholds 0.0). Accuracy of segmentation was visually checked by two authors (YB and MD). Discrepancies in the analyzed fields of view were solved by manual tracing after *ad hoc* adjustment of brightness and contrast. The dataset of cell metrics is available as Supplementary information.

## Results

The clinical characteristics of the patients are indicated in Table 1. Figure 1 and supplementary video 1 shows the case of patient 1 which had multiple scars from panretinal photocoagulation for proliferative diabetic retinopathy. The successive color fundus photographs taken over 44 months after laser showed progressive enlargement of scars. Time-lapse imaging of a scar in the posterior pole showed the presence of two hyperpigmented spots along the edges, which moved in opposite directions during expansion (Figure 1, B-E). While remaining within the edge of the scar during the entire follow-up, these spots elongated as the scar enlarged (Figure 1E). Another example of mobilization of melanocytic spot is shown in supplementary video 2, in which two hyperpigmented spot in the edges of atrophy are seen moving in opposite directions. Patient 2 (figure 2 and supplementary video 3) showed extensive laser scars of various size in the in posterior pole. Time-lapse imaging of infrared SLO images recapitulating 5 years of evolution showed the expansion of a large scar with two smaller scars apparently moving away from it (Figure 2B and 2C). This was interpreted as an outward displacement of the RPE strip between the scars.

**Figure 1.**
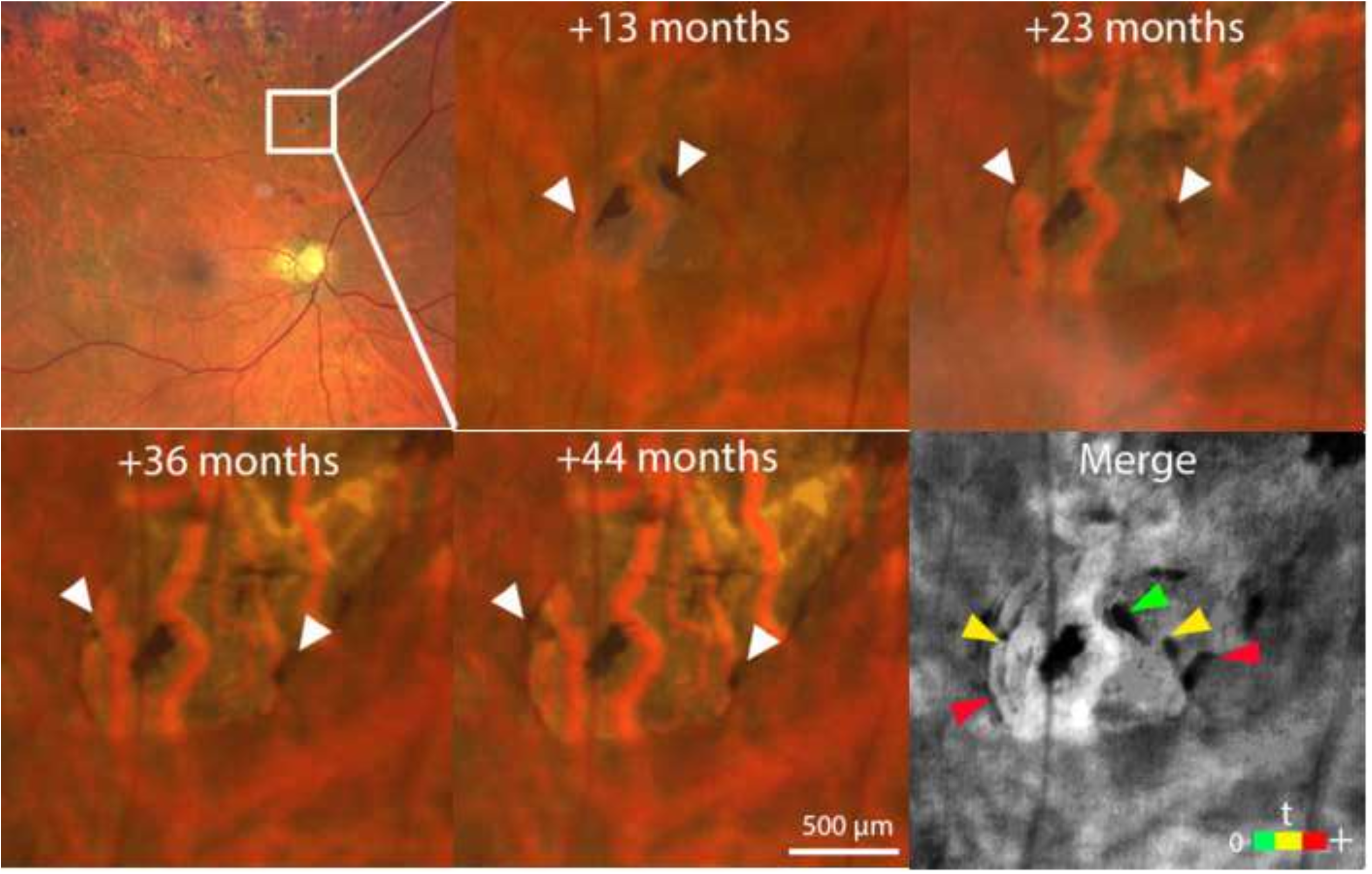
Case 1. A, color fundus photograph showing panretinal photocoagulation in the superior retina that caused multiple scars. B to E, evolution over time of the scar in the squared area showing progressive enlargement over 2.6 years (i.e. up to 44 months after laser); Arrowheads point to two hyperpigmented foci on the edges of the scar that are displaced in opposite directions. Note also the progressive elongation of the hyperpigmentation. F, merged image. Time indicates delay from laser; the color bar indicates chronology. See also supplementary video 1.

**Figure 2.**
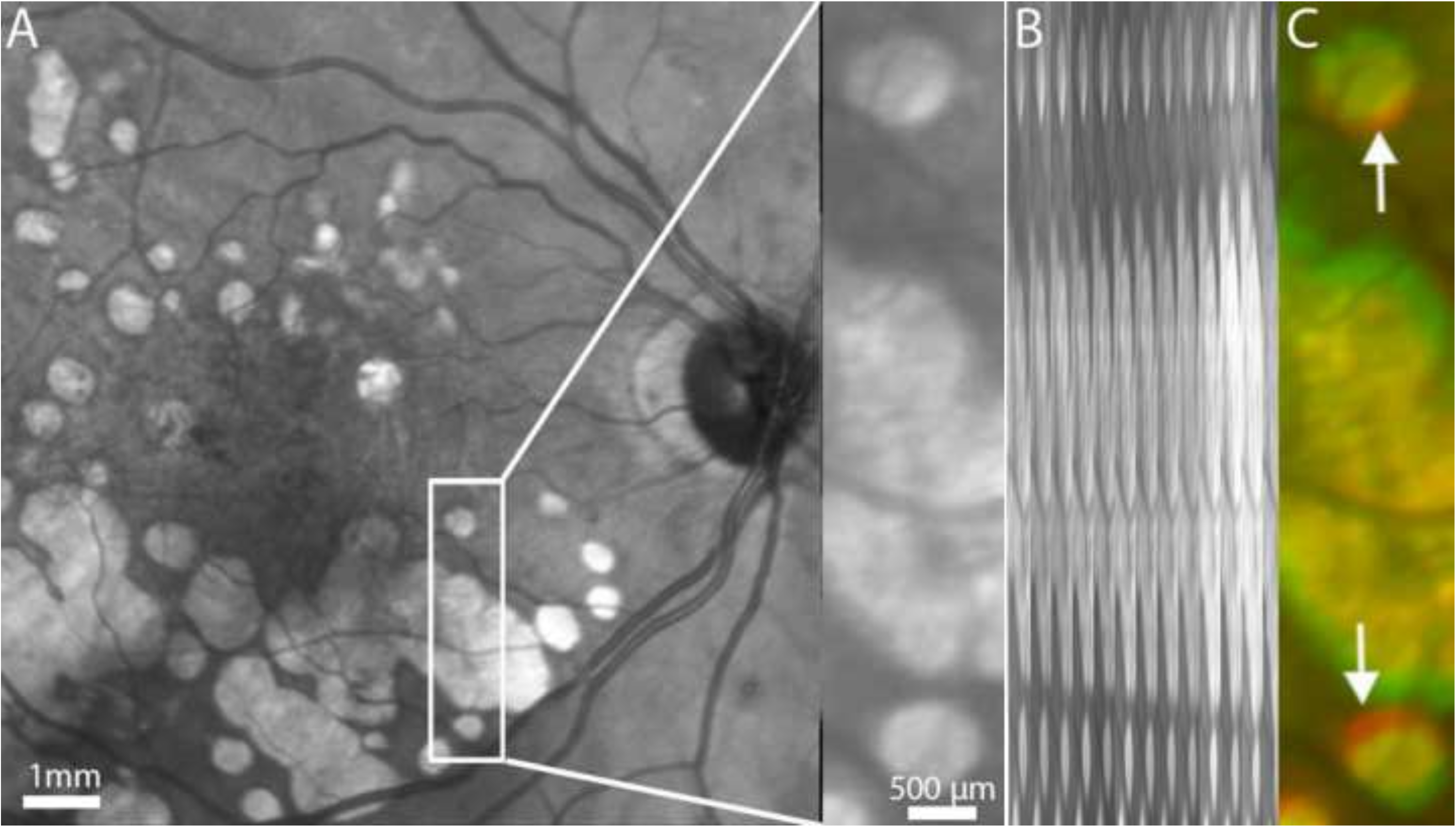
Case 2. A, scanning laser ophthalmoscopy (SLO) infrared imaging. B, kinetogram of the squared area showing progressive enlargement of a scar over 5 years. C, color coded merge of the first and last images. The red rims (arrows) indicate displacement of the thin RPE band between two scars. See also supplementary video 3.

In addition to conventional imaging, cases 3, 4 and 5 were prospectively imaged by TFI AOO in an attempt to track individual RPE cells (figures 3 and 4, and supplementary videos 4 to 6). In all cases, TFI AOO visualized a mosaic of dark spots in the margin of the scars, with a density ranging from 1280 to 3254 /mm². Time-lapse sequences of TFI images revealed that there were movements of this mosaic during expansion of scars. The movements of XXX dark spots were tracked, and their direction reported on directional maps. These movements were detected up to several hundred μm away from the edges of scars. In a given scar, the direction of movements varied, that is, cells were not only moving radially away from the scar. In case 3, in a strip of RPE between two scars, RPE cells were moving tangentially in opposite directions (Fig. 3D and supplementary video 4). In case 4, movements were detected in the nasal and temporal sides of scars but were less important along the superior and inferior sides (Figure 4B and supplementary video 5). Case 5 showed a variety of directions of movements, with tangent and even retrograde movements (figure 4D and supplementary video 6).

**Figure 3.**
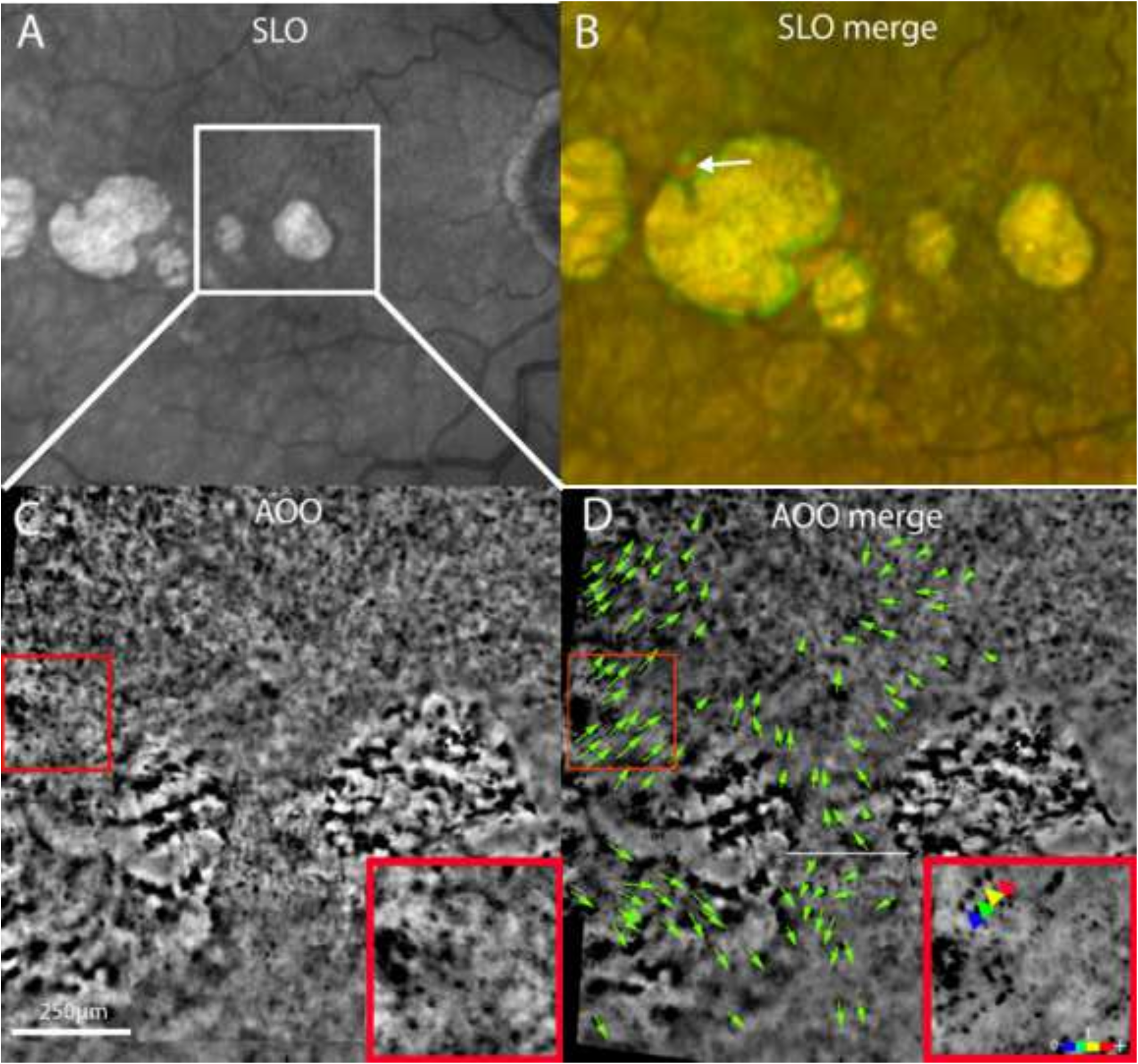
Case 3. A, SLO infrared images. B, red-green color-coded SLO image of images taken 2 years apart; the green rim indicates enlargement of the scar. Note the displacement of a hyperpigmented spot (arrow; the green spot indicates the final position; this melanocytic spot is shown in supplementary video 2). C, TFI AOO image. Note the irregular mosaic of dark spots, presumably RPE cells. In the merged image (D), the green arrows indicate the displacement of individual dark spots. Note the opposite movements within the RPE strip between two scars in the center of the image, the white line figuring an apparent demarcation line. The insert shows successive positions of a single spot; chronological order is indicated in the color bar (See also supplementary video 4).

**Figure 4:**
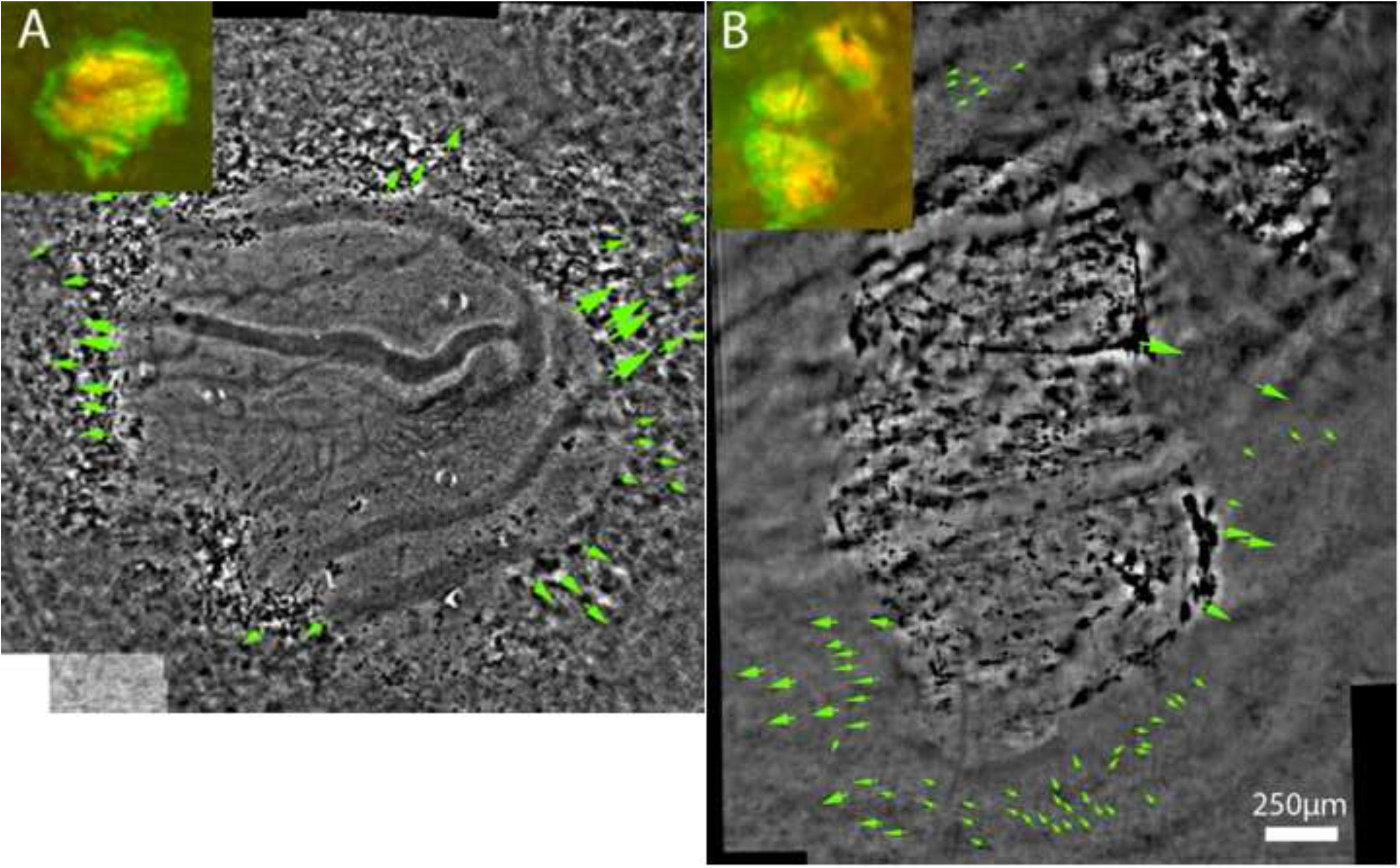
Infrared and color-coded merged fundus images and TFI AOO images recapitulating movements of a mosaic of dark spots in cases 4 (A and C; follow-up 14 months) and 5 (B and D; follow-up 13 months). Inserts in A and B are SLO color-coded merged images; the green rim shows progression of atrophy. The direction of arrows indicates the movements during follow-up in C and D. Note the variable orientation of the movement vectors; in the RPE strip between the two central scars, movements are tangent to scar edges and in opposite directions. See also supplementary videos 5 and 6.

To document the phenotype of individual RPE cells around laser scars, a RPE-choroidal flat-mount from a donor eye with a history of sectorial scatter laser photocoagulation was examined by confocal microscopy. The optimal observation of the RPE on flat-mounts required to ablate the retina; this could potentially cause damage to the RPE since it is known that an atrophic retina adheres to the Bruch membrane. Therefore, prior to removal of the retina, *ex vivo* NIRAF imaging was performed which delineated the edges of the areas lacking RPE (Fig. 5A). This shows post-laser confluence of scars, demonstrating that there has been expansion of scars after photocoagulation. Immediately following tissue processing, comparison of pre-ablation images with confocal microscopy images allowed to verify that the edges of scars were minimally altered by retinal ablation (compare figures 4A and 5). Hence, we considered that the RPE cells lining the atrophy, that is, RPE cells within the progression front, were minimally affected by tissue processing. Confocal microscopy of phalloidin-labelled RPE cells, which reveals actin fibers, showed that the RPE population was heterogeneous both in size and shape. All cells were RPE65+ (not shown). Cells along edges of scars were often elongated, with their long axis tangent to the edge of the atrophy (Figure 5). Figure 5A shows that clusters of enlarged RPE cells were distributed between atrophic scars, possibility corresponding to healed laser scars [21,22]. The edges of scars often showed a linear actin labelling forming a multicellular structure along the first line of RPE cells along the atrophic zone, and extending over several hundred microns. Actin cables did not entirely surround scars, instead showing multiple interruptions (figure 6C). Enlarged and elongated cells formed fascicles in strips of RPE between scars (Figure 6D and E).

**Figure 5.**
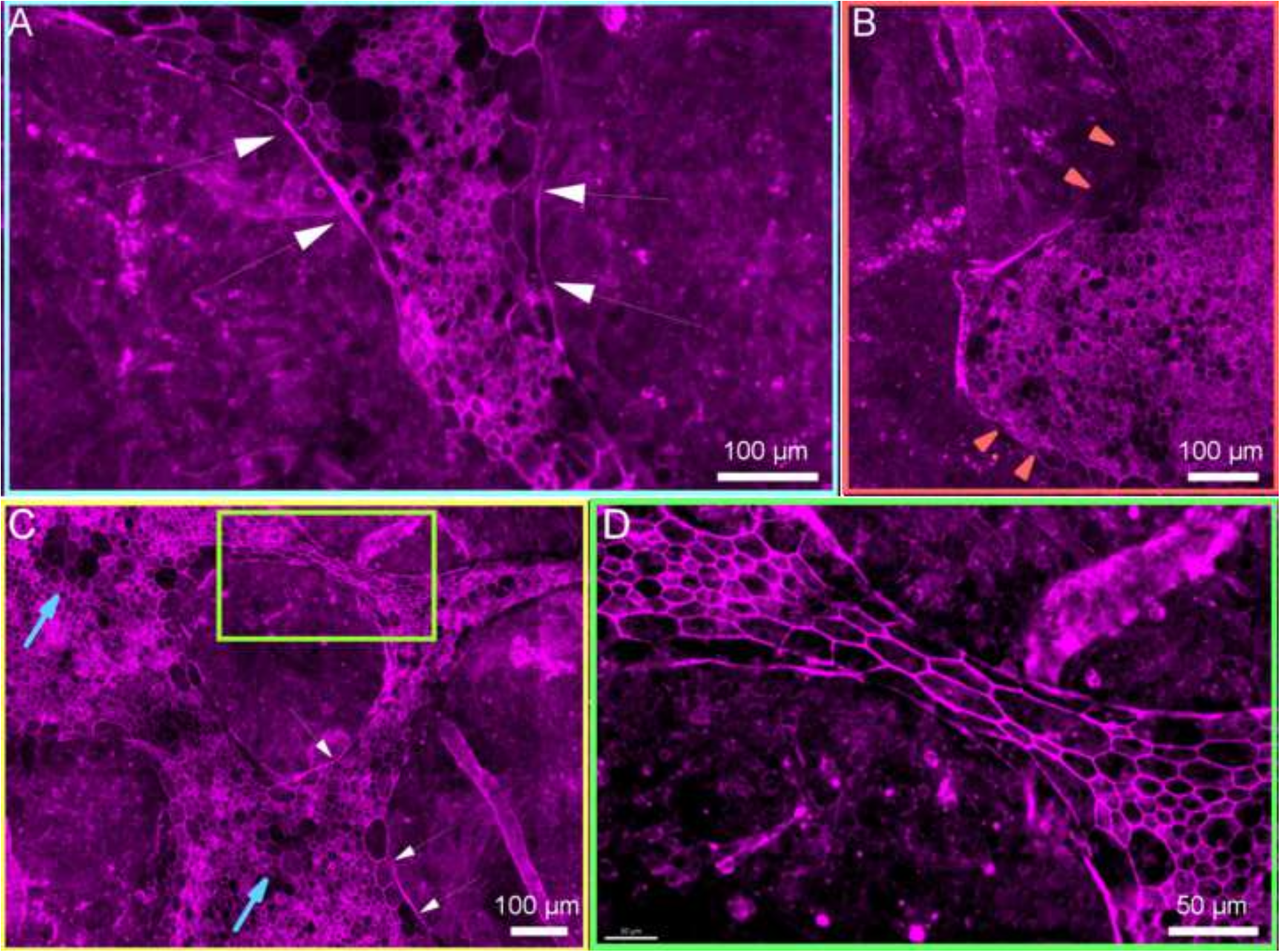
Histology examination of a human eye with laser scars (84-year-old female). A, *ex vivo* NIRAF SLO imaging prior to histology processing. The optic nerve is indicated by the circle and the fovea by the arrowhead. The colored squares indicate the location of subsequent panels. Comparison with panel B demonstrates the general preservation of the edges of the scars after removal of the retina. B to E, mapping of the metrics of RPE cells (. B and D: cell size; C and E: aspect ratio). B and C, large field views; D and E, magnification of the area shown in the white squares. Note that elongated cells are mostly along margins, while enlarged cells are either along the edges or in cluster (example shown by the blue arrows in D).

**Figure 6.**
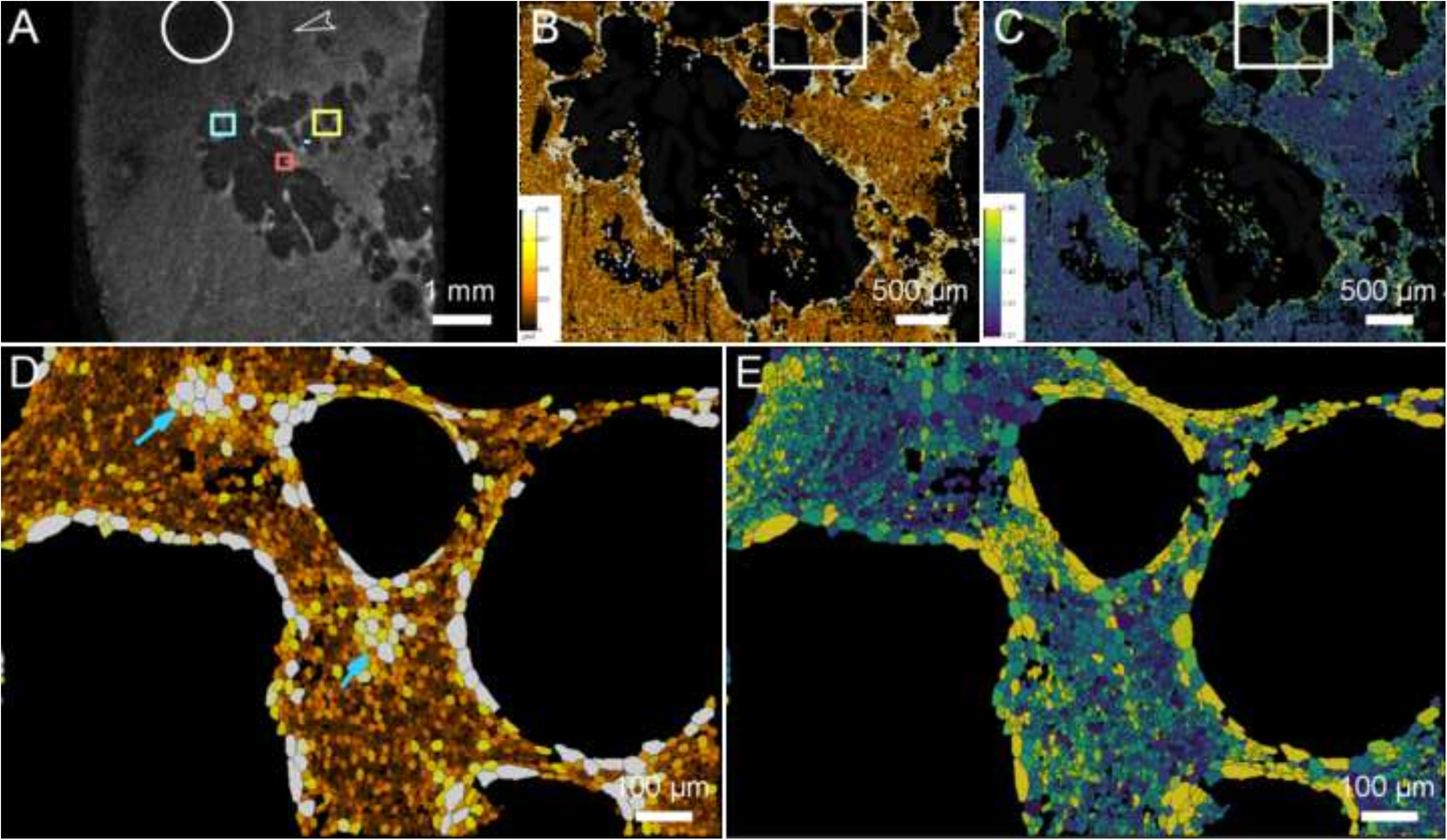
Confocal microscopy of phalloidin-labelled (magenta) RPE cells. In A, note the presence of segments of phalloidin-affine structures along the edge of the scar (white arrowheads), and the heterogeneity of cell size. In B, note the interruption of the actin cable (red arrowheads) along the edge of a scar. C and D, successive magnifications of a strip of RPE cells between two scars. In C, blue arrows point to a cluster of enlarged RPE cells, possibly representing healed scars; similar sized RPE cells are also present along the edges of scars (white arrowheads). D shows a fascicle of parallel, elongated RPE cells. Note that cell elongation is accentuated in the center of the fascicle.

## Discussion

It has been well established that epithelial biomechanics are crucially involved in closure of wounds in postmitotic epithelia from a variety of species and tissues; their possible implication in wound expansion as well has been documented more recently [10]. The finding that cell contractility may also contribute to hole expansion *in vitro* demonstrated that the net effect of cell contractility on epithelial defects results from a balance of supracellular organized inward forces (tending to close holes by the action of actin cables) versus outward forces (cell contractility) which tends to enlarge holes. It is known that the capability of the actin cable to close an epithelial defect depends on the radius of the hole; it is less efficient in larger holes [1]. Hence, this balance may be altered by factors such as the size of defects, cell density or adhesion to the substrate. Notably, these processes have not been described *in vivo* in humans.

Here, in human eyes with laser scars in the RPE we searched for manifestations of epithelial biomechanics by multimodal imaging and confocal microscopy. In patients, multimodal time-lapse imaging revealed that expanding scars were accompanied by dynamic changes in the surrounding RPE whose kinetics closely matched the progression of atrophy. In particular, high-resolution imaging suggested collective movements of a mosaic that closely resembled that reported for the RPE cells. These displacements were spatially and temporally correlated to the enlargement of scars.

This raises the question of the possible mechanisms of such movements. By histology, we found numerous elongated RPE cells, often lined along the edges of scars. Of particular interest was the finding of fascicles of elongated cells between adjacent scars. Such stretching has been reported *in vitro* in strips of epithelial cells between adjacent holes [10], and this precedes epithelial fracture and subsequent fusion of adjacent holes. Furthermore, we also found numerous stress fibers located in RPE cells along the edges of atrophy, forming segments of actin cables. These actin cables were associated with regularization of the edges of the scars and most often were convex relative to the scar, which is seen in epithelial holes undergoing a purse string effect. We also found clusters of enlarged RPE cells; accordingly, an *in vitro* study found that wound healing of human RPE cells could lead to the development of enlarged RPE cells surrounded by radially elongated cells [22]. A similar process of tissue repair causing enlarged cells in post-mitotic epithelia has been described as compensatory cell hypertrophy, which is another pathway for the restoration of epithelial coverage [23]. Compensatory cell hypertrophy occurs at the cost of increasing tissue tension, hence further aggravating biomechanical stress. Taken together, these stigmatas of long-lasting biomechanical traction support the presence of chronically increased epithelial biomechanical stress. Hence, we hypothesize that in eyes with expanding laser scars, RPE cells are outwardly dragged, similarly to a carpet dragging effect. This occurs despite the presence of actin cables, which are counteracting the outward dragging; The large size of the atrophy and the fragmentation of actin cables are probably weakening the purse-string effect, hence lessening inwardly oriented force vectors therefore shifting the balance of forces toward enlargement of the holes. However, limits to these conclusions are based on the fact the longitudinal imaging observations and the histological findings were obtained from different individuals, and the histological analysis represents a static examination of a single donor eye.

Other limits of our study are the small sample of patients and the indirect visualization of RPE cells. Indeed, the correspondence of the mosaic of dark spots seen by AOO with that of RPE cells is uncertain, because the density of dark spots detected by TFI was somewhat inferior to the reported density of human RPE cells seen by TFI [18]. This however does not exclude the possibility that this mosaic represented a subset of RPE cells. Hence, it is reasonable to assume that the mosaic detected by TFI was a surrogate of the overall movements of the RPE. Although the presence of elongated cells, stress fibers, and actin-cable-like structures is consistent with a tractional phenotype, these observations do not directly demonstrate contractility, the direction or magnitude of mechanical forces, or a causal role of these structures in scar enlargement. Furthermore, as for most chronic pathological processes, it is likely that several pathways act synergistically, therefore, our data does not exclude the implication of other mechanisms such as epithelial-mesenchymal transition, cell degeneration, oxidative stress and/or cell death. Another limit of our study is that we could not map the expression of the catalog of antigens involved in biomechanical tractions, because the use of flat-mounts limits the number of antigens to the number of wavelengths of the confocal microscope. Other elements of the healing process such as lamellipodia, which are cellular extension driving cell migration [24] also remains to be documented to acquire a full picture of the activation of healing-related mechanisms in the RPE.

To sum up, around expanding RPE scars we detected dynamic and histological features that are reminiscent of canonical manifestations of biomechanical stress. To our knowledge, this is the first report suggesting the implication of biomechanical traction in an epithelial disease in humans. This supports the hypothesis that collective movements of RPE cells elicited by the tissue-scale contraction of stress fibers contributed to the enlargement of laser scars. The conceptual framework of epithelial biomechanics physiopathology, a research field at the interface of physics and biology, may therefore help to better understand the expansion of RPE atrophy and to explore the role of other factors influencing biomechanical stress.

Other types of RPE atrophies are also known to expand such as GA [25], severe myopia [26] and RPE dystrophies [27]. In the case of GA, we have shown by time-lapse imaging that hyperpigmented clumps located in the RPE in the margins of lesions may move ahead of atrophy progression [28]. Therefore, documenting biomechanical stress in the RPE may be of interest in these diseases as well.

## Supplementary videos

**Supplementary video 1**. Case 1. Time-lapse videosequence of color fundus photographs. The red component of the color image was filtered to highlight the contrast of melanin (field of view 8 mm x 8 mm; duration in real time 20 months; 2fps). In the square, note the outward displacement of the pigmented spots located in margins of the scar (arrows).

**Supplementary video 2**. Case 3. Time-lapse videosequence of near infrared autofluorescence SLO images of the eye shown in figure 3. Note the outward displacement of hyperfluorescent spots (arrows) (field of view 9 mm x 9 mm; duration in real time 58 months; 2fps).

**Supplementary video 3**. Case 2. Time-lapse videosequence of infrared SLO images of the eye shown in figure 2 (field of view 9 mm x 9 mm; duration in real time 60 months; 2fps). Note the displacement of satellite scars (arrows) during the enlargement of the larger scar between them.

**Supplementary video 4**. Case 3. Time-lapse videosequence of TFI AOO images of the eye shown in figure 3. Note the outward displacement of the mosaic of dark spots (arrows) (field of view 5mmx5mm; duration in real time 16 months; 2fps).

**Supplementary video 5**. Case 4. Time-lapse videosequence of TFI AOO images of the eye shown in figure 4A (field of view 2mmx2mm; duration in real time 14 months; 2fps). In the squares, note the outward displacement of the mosaic of dark spots (arrows).

**Supplementary video 6**. Case 5. Time-lapse videosequence of TFI AOO images of the eye shown in figure 4B (field of view 2mmx3mm; duration in real time 13 months; 2fps). In the squares, note the outward displacement of the mosaic of dark spots.

## References

[1] Ladoux B, Mège R-M. Mechanobiology of collective cell behaviours. Nat Rev Mol Cell Biol 2017;18:743–57. 10.1038/nrm.2017.98.

[2] Bement WM, Forscher P, Mooseker MS. A novel cytoskeletal structure involved in purse string wound closure and cell polarity maintenance. J Cell Biol 1993;121:565–78. 10.1083/jcb.121.3.565.

[3] Martin P, Lewis J. Actin cables and epidermal movement in embryonic wound healing. Nature 1992;360:179–83. 10.1038/360179a0.

[4] Movrin V, Krajnc M. Initiation of epithelial wound closure by an active instability at the purse string. Biophys J 2025;124:107–14. 10.1016/j.bpj.2024.11.008.

[5] Roffay C, Chan CJ, Guirao B, Hiiragi T, Graner F. Inferring cell junction tension and pressure from cell geometry. Development 2021;148:dev192773. 10.1242/dev.192773.

[6] Shen Y, O’Byrne J, Schoenit A, Maitra A, Mège R-M, Voituriez R, et al. Flocking and giant fluctuations in epithelial active solids. Proceedings of the National Academy of Sciences 2025;122:e2421327122. 10.1073/pnas.2421327122.

[7] Fernandez-Gonzalez R, Harris TJC. Contractile and expansive actin networks in Drosophila: Developmental cell biology controlled by network polarization and higher-order interactions. Curr Top Dev Biol 2023;154:99–129. 10.1016/bs.ctdb.2023.02.005.

[8] Brock J, Midwinter K, Lewis J, Martin P. Healing of incisional wounds in the embryonic chick wing bud: characterization of the actin purse-string and demonstration of a requirement for Rho activation. J Cell Biol 1996;135:1097–107. 10.1083/jcb.135.4.1097.

[9] Kamran Z, Zellner K, Kyriazes H, Kraus CM, Reynier JB, Malamy JE. In vivo imaging of epithelial wound healing in the cnidarian Clytia hemisphaerica demonstrates early evolution of purse string and cell crawling closure mechanisms. BMC Dev Biol. 2017 Dec 19;17(1):17.

[10] Lv J-Q, Chen P-C, Chen Y-P, Liu H-Y, Wang S-D, Bai J, et al. Active hole formation in epithelioid tissues. Nat Phys 2024;20:1313–23. 10.1038/s41567-024-02504-1.

[11] Sparrow JR, Hicks D, Hamel CP. The Retinal Pigment Epithelium in Health and Disease. Curr Mol Med 2010;10:802–23. 10.2174/156652410793937813.

[12] Brancato R, Pece A, Avanza P, Radrizzani E. Photocoagulation scar expansion after laser therapy for choroidal neovascularization in degenerative myopia. Retina 1990;10:239–43. 10.1097/00006982-199010000-00002.

[13] Lee TC, Lee S-W, Dinkin MJ, Ober MD, Beaverson KL, Abramson DH. Chorioretinal scar growth after 810-nanometer laser treatment for retinoblastoma. Ophthalmology 2004;111:992–6. 10.1016/j.ophtha.2003.08.036.

[14] Maeshima K, Utsugi-Sutoh N, Otani T, Kishi S. Progressive enlargement of scattered photocoagulation scars in diabetic retinopathy. Retina 2004;24:507–11. 10.1097/00006982-200408000-00002.

[15] Sutoh N, Muraoka K, Tokui K, Takahashi K. Wide-angle indocyanine green angiography in diabetic retinopathy after panretinal photocoagulation. Japanese Journal of Ophthalmology 1995;49:121–7.

[16] Tarau I-S, Berlin A, Curcio CA, Ach T. The Cytoskeleton of the Retinal Pigment Epithelium: from Normal Aging to Age-Related Macular Degeneration. Int J Mol Sci 2019;20:E3578. 10.3390/ijms20143578.

[17] Chan CM, Huang DY, Sekar P, Hsu SH, Lin WW. Reactive oxygen species-dependent mitochondrial dynamics and autophagy confer protective effects in retinal pigment epithelial cells against sodium iodate-induced cell death. J Biomed Sci. 2019 May 22;26(1):40.

[18] Laforest T, Künzi M, Kowalczuk L, Carpentras D, Behar-Cohen F, Moser C. Transscleral Optical Phase Imaging of the Human Retina. Nat Photonics 2020;14:439–45. 10.1038/s41566-020-0608-y.

[19] Gofas-Salas E, Lee DMW, Rondeau C, Grieve K, Rossi EA, Paques M, et al. Comparison between Two Adaptive Optics Methods for Imaging of Individual Retinal Pigmented Epithelial Cells. Diagnostics (Basel) 2024;14:768. 10.3390/diagnostics14070768.

[20] Stringer C, Wang T, Michaelos M, Pachitariu M. Cellpose: a generalist algorithm for cellular segmentation. Nat Methods 2021;18:100–6. 10.1038/s41592-020-01018-x.

[21] Ach T, Huisingh C, McGwin G Jr, Messinger JD, Zhang T, Bentley MJ, Gutierrez DB, Ablonczy Z, Smith RT, Sloan KR, Curcio CA. Quantitative autofluorescence and cell density maps of the human retinal pigment epithelium. Invest Ophthalmol Vis Sci. 2014 Jul 17;55(8):4832–41. doi: 10.1167/iovs.14-14802. PMID: 25034602; PMCID: PMC4123894.

[22] Bailey-Steinitz LJ, Shih Y-H, Radeke MJ, Coffey PJ. An in vitro model of chronic wounding and its implication for age-related macular degeneration. PLoS ONE 2020;15:e0236298. 10.1371/journal.pone.0236298.

[23] Tamori Y, Deng WM. Compensatory cellular hypertrophy: the other strategy for tissue homeostasis. Trends Cell Biol. 2014 Apr;24(4):230–7. doi: 10.1016/j.tcb.2013.10.005. Epub 2013 Nov 14. PMID: 24239163; PMCID: PMC4022146.

[24] Anon E, Serra-Picamal X, Hersen P, Gauthier NC, Sheetz MP, Trepat X, et al. Cell crawling mediates collective cell migration to close undamaged epithelial gaps. Proceedings of the National Academy of Sciences 2012;109:10891–6. 10.1073/pnas.1117814109.

[25] Shen LL, Sun M, Khetpal S, Grossetta Nardini HK, Del Priore LV. Topographic Variation of the Growth Rate of Geographic Atrophy in Nonexudative Age-Related Macular Degeneration: A Systematic Review and Meta-analysis. Invest Ophthalmol Vis Sci. 2020 Jan 23;61(1):2. doi: 10.1167/iovs.61.1.2. PMID: 31995152; PMCID: PMC7205189.

[26] Wender JD, Johnson RN, McDonald HR, Jumper JM, Fu AD, Lujan BJ, et al. Continued vision loss from progressive expansion of inactive chorioretinal scars in a myopic patient with well-controlled sympathetic ophthalmia. Retin Cases Brief Rep 2012;6:383–7. 10.1097/ICB.0b013e31823f09bb.

[27] Bommakanti N, Young BK, Sisk RA, Berrocal AM, Duncan JL, Bakall B, et al. Classification and Growth Rate of Chorioretinal Atrophy after Voretigene Neparvovec-Rzyl for RPE65-Mediated Retinal Degeneration. Ophthalmol Retina 2024;8:42–8. 10.1016/j.oret.2023.08.017.

[28] Paques M, Norberg N, Chaumette C, Sennlaub F, Rossi E, Borella Y, Grieve K. Long Term Time-Lapse Imaging of Geographic Atrophy: A Pilot Study. Front Med (Lausanne). 2022 Jun 22;9:868163. doi: 10.3389/fmed.2022.868163. PMID: 35814763; PMCID: PMC9257004.

